# Viral and host small RNA transcriptome analysis of SARS-CoV-1 and SARS-CoV-2-infected human cells reveals novel viral short RNAs

**DOI:** 10.1101/2022.12.27.522023

**Authors:** Tom Driedonks, Lyle H. Nyberg, Abigail Conte, Zexu Ma, Andrew Pekosz, Eduard Duban, Holger Sültmann, Andrey Turchinovich, Kenneth Witwer

## Abstract

RNA viruses have been shown to express various short RNAs, some of which have regulatory roles during replication, transcription, and translation of viral genomes. However, short viral RNAs (svRNAs) generated by SARS-CoV-1 and SARS-CoV-2 remained largely unexplored, mainly due limitations of the widely used library preparation methods for small RNA deep sequencing and corresponding data processing. By analyzing publicly available small RNA-seq datasets, we observed that human cells infected by SARS-CoV-1 or SARS-CoV-2 produce multiple short viral RNAs (svRNAs), ranging in size from 15 to 26 nt and deriving predominantly from (+) RNA strands. In addition, we verified the presence of the five most abundant SARS-CoV-2 svRNAs in SARS-CoV-2-infected human lung adenocarcinoma cells by qPCR. Interestingly, the copy number of the observed SARS-CoV-2 svRNAs dramatically exceeded the expression of previously reported viral miRNAs in the same cells. We hypothesize that the reported SARS-CoV-2 svRNAs could serve as biomarkers for early infection stages due to their high abundance. Finally, we found that both SARS-CoV-1 and SARS-CoV-2 infection induced up- and down-regulation of multiple endogenous human short RNAs that align predominantly to protein-coding and lncRNA transcripts. Interestingly, a significant proportion of short RNAs derived from full-length viral genomes also aligned to various hg38 sequences, suggesting opportunities to investigate regulatory roles of svRNAs during infection. Further characterization of the small RNA landscape of both viral and host genomes is clearly warranted to improve our understanding of molecular events related to infection and to design more efficient strategies for therapeutic interventions as well as early diagnosis.

## INTRODUCTION

The 2019 outbreak of SARS-CoV-2, the causative agent of coronavirus disease COVID-19 [Lu et al, 2020; Hu et al, 2021], led to more than six million deaths by the year 2022 [World Health Organization, 2022]. Amidst unprecedented mobilization of resources for diagnosis and treatment, it remains crucial to develop a more detailed understanding of the molecular perturbations accompanying SARS-CoV-2 infection. Most transcriptomic studies of infection with SARS-CoV-2 and related viruses focus on long RNA biotypes, with limited but highly interesting attention to small RNAs. Precise characterization of the host and viral small RNA landscape in infected cells could provide important insights. Small RNA-related molecular mechanisms that regulate the viral life cycle and could be targeted for therapeutic interventions. Meanwhile, the relative stability of small RNAs could enable novel diagnostic and prognostic tools, for example, to detect the presence of SARS-CoV-2 in biological fluids and mucosal surfaces for “as-early-as-possible” diagnosis or detection of low-level virus persistence.

Small non-coding RNAs (sncRNAs) are known to mediate diverse regulatory functions during cell proliferation, differentiation, apoptosis and response to infection [Zhang et al, 2021]. MicroRNAs, piwi-interacting RNAs and tRNA-derived RNA fragments are some examples of the most widely studied sncRNAs. Changes in host microRNA expression accompanied by subsequent deregulation of microRNA target genes have been previously observed in cells infected with different viruses including SARS-CoV-1 and SARS-CoV-2 [Girardi et al, 2018; Barbu et al, 2020, Wyler et al, 2021]. However, differential expression of other small RNA species has not been as well addressed, presumably due to the fact that widely used protocols for small RNA sequencing library preparation preferentially incorporate RNAs containing 5’-Phosphate and 3’-OH groups. Viruses may also encode their own sncRNAs to regulate replication, transcription, and translation of viral genomes as well as to influence immune evasion and inflammatory response.

In this work, we characterized small RNA expression perturbations caused by SARS-CoV-1 and SARS-CoV-2 infection in the lung cell line Calu-3 using publicly available datasets obtained using ligation-independent library preparation techniques [Wyler et al, 2021]. Unlike in the original publication, where small RNA-seq data analysis was limited to microRNAs, we applied a mapping pipeline to account for all human RNA biotypes. Our analysis confirmed remarkably different responses of small RNA expression during SARS-CoV-1 and SARS-CoV-2 infection in Calu-3 cells. Thus, SARS-CoV-2 induced differential expression of a dramatically larger number of small RNAs as compared with SARS-CoV-1. Importantly, these small RNA fragments were aligned predominantly to protein-coding and lncRNA genes. At the same time, despite robust changes in small RNA expression, only a few miRNAs were affected, suggesting a limited role of host miRNAs in the progression of infection. The mechanisms of formation and biological roles of the aforementioned short RNAs aligned to mRNAs and lncRNAs remains to be elucidated.

Finally, we show that SARS-CoV-1 and SARS-CoV-2 infected cells contain a variety of short viral RNAs (svRNAs) which have dramatically higher expression as compared with previously identified viral miRNAs. Interestingly, a significant percentage of the identified svRNAs also aligned to various locations in the human genome with zero-mismatch tolerance.

While further studies are required to elucidate the mechanisms of biogenesis and biological role of these svRNAs, it is plausible that SARS-CoV-2 derived svRNAs might serve as ultrasensitive novel biomarkers for early diagnosis or detection of persistent infection. Our work provides an update to small RNA expression during SARS-CoV-2 infection and suggests several independently validated virus-derived short RNA fragments as possible markers of early or low-level, persistent infection.

## MATERIALS AND METHODS

### Raw small RNA-seq datasets

Raw fastq files associated with a previous publication [Wyler et al, 2021] were downloaded from the GEO database (accession number GSE148729). The small RNA sequencing dataset included six controls (SRR11550015, SRR11550016, SRR11550017, SRR11550018, SRR11550031, SRR11550032) and twelve samples from cells infected with either SARS-CoV-1 (SRR11550019, SRR11550020, SRR11550021, SRR11550022, SRR11550023, SRR11550024) or SARS-CoV-2 (SRR11550025, SRR11550026, SRR11550027, SRR11550028, SRR11550029, SRR11550030). The total RNA sequencing and mRNA sequencing datasets from the same mock and SARS-CoV-1/2 infected cells were used to evaluate the origin of certain small RNA-seq reads.

### Bioinformatics analysis

Raw small RNA-seq reads were trimmed of poly-A tails and adapter sequences including the first 3 nucleotides using cutadapt software (*cutadapt -u 3 input.fastq* | *cutadapt -a AAAAAAAA -o output.fastq*). Reads shorter than 15 nucleotides were discarded. The trimmed and size-selected reads were further aligned to either SARS-CoV-1 or SARS-CoV-2 reference genomes (NC_004718.3 and NC_045512.2, respectively) with *bowtie,* allowing no mismatches (- *v 0* option). The reads derived from forward (+) and reverse (-) strands were differentiated using --*norc* and --*nofw* parameters, respectively, during alignment with *bowtie*. Reads exclusively aligned to either SARS-CoV-1 or SARS-CoV-2 were extracted using a combination of *samtools*, *bam2fastx* and *bowtie* packages. Sorted and indexed BAM files were generated using *samtools,* and the alignments to the reference viral genomes were visualized in *IGV*. Small RNA-seq datasets (filtered from all SARS-CoV-1/2 reads) were mapped to human references in a sequential manner; specifically, trimmed and size-selected reads were first mapped to RNA species having low sequence complexity and/or a large number of repeats, including rRNA, tRNA, RN7S, snRNA, snoRNA/scaRNA, vault RNA, RNY (custom-curated references from NCBI RefSeq and GENCODE) and the mitochondrial chromosome, from GRCh38. All reads that did not map to the aforementioned RNAs were sequentially aligned to mature miRNA (miRBase 22 release), pre-miRNA (miRBase 22 release), protein-coding mRNA transcripts and long non-coding RNAs (custom-curated references from GENCODE v32). Remaining unmapped reads were aligned to the remaining transcriptome (GENCODE v32), containing mostly pseudogenes and non-protein-coding parts of mRNA transcripts. Finally, reads not mapping to the human transcriptome were aligned to the human primary GRCh38 assembly, corresponding to reads derived from introns and intergenic regions. The numbers of reads mapped to each RNA reference type were extracted using eXpress software based on a previous publication [Roberts and Pachter, 2013]. The differential gene expression analysis of both small RNA-seq and mRNA-seq was done using R/Bioconductor packages edgeR and limma as described previously [Law et al, 2016].

### RT-qPCR validation of svRNA from independent infection experiments

Human lung adenocarcinoma (A549) cells overexpressing the ACE2 receptor were seeded at 1 million cells per well in a 6-well plate. Three wells were infected with at a multiplicity of infection (MOI) of 5.0 tissue culture infectious units (TCID50) per cell with SARS-CoV-2/USA/DC-HP00007/2020 (GenBank: MT509464.1) virus, and three wells remained uninfected. Infections were performed as previously described [Park et al, 2022]. RNA was isolated from cells 24h after infection using the miRNeasy micro kit (Qiagen, Hilden, Germany) and Zymo-Spin IIICG columns. RNA was eluted in 50 μl RNase-free water, and RNA concentration was measured by NanoDrop One UV-Vis spectrophotometer (ThermoFisher). cDNA was prepared by miScript RT II cDNA kit (Qiagen) according to manufacturer’s instructions, using HiFlex buffer and 4 μg RNA input. No-RT controls were included to confirm the absence of co-isolated cellular DNA. Next, cDNA was diluted tenfold and used in qPCR; 2 μl of diluted cDNA template was mixed with 4 μl QuantiTect SYBR Green PCR Master Mix (Qiagen), 100 nM forward primers (see **Table 1** for sequences) and 100 nM miScript universal reverse primer. After an initial polymerase activation step (15 minutes at 95C), 50 cycles of the following program were run: 15 sec at 94C, 30 sec at 55C, 30 sec at 70C, followed by melting curve analysis on a BioRad CFX96 machine.

**Table 1.**
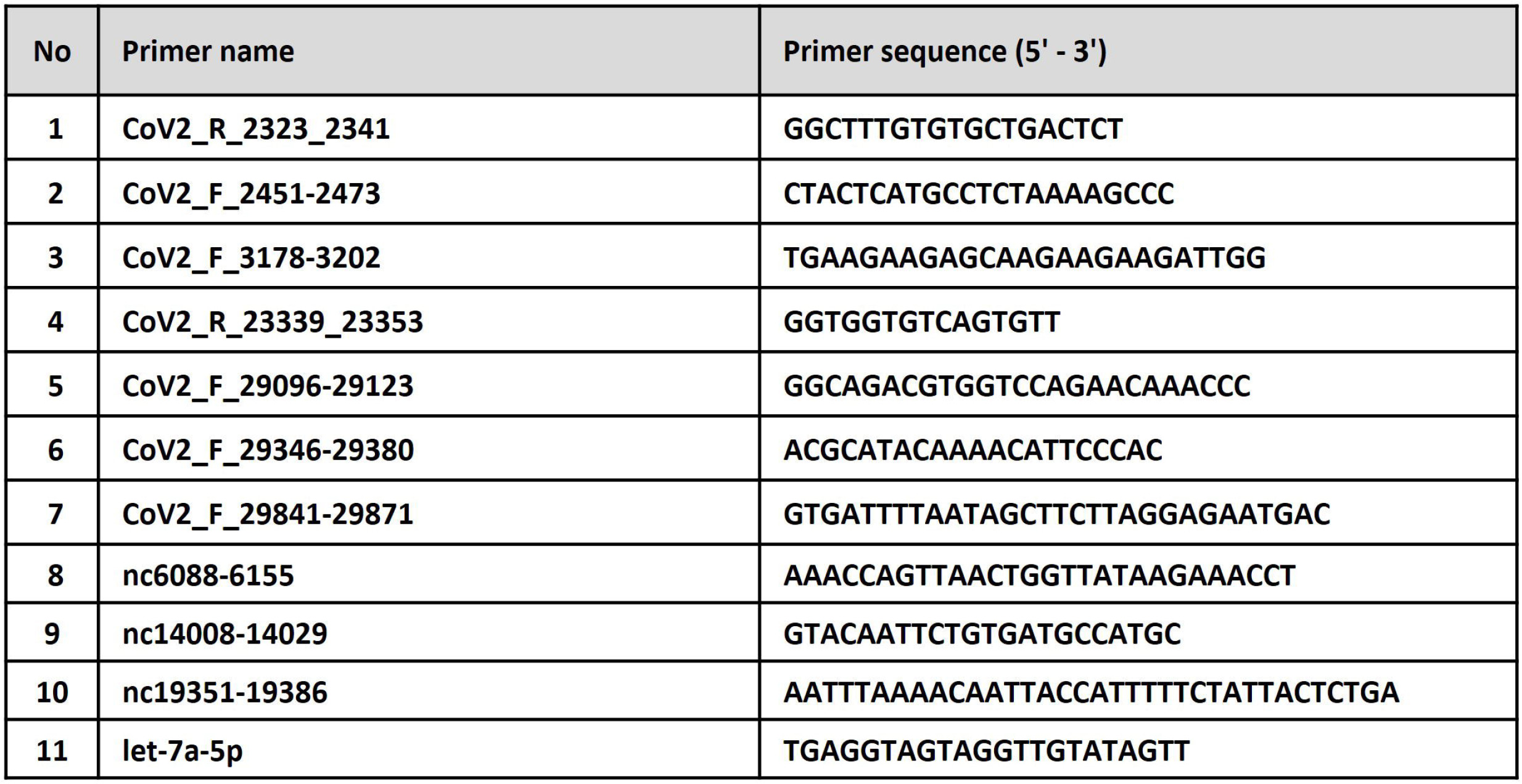
RT-qPCR primer sequences. Numbers in primers’ names represent the start and stop sites of the sequences in the SARS-CoV-2 genome (reference sequence NC_045512.2). Primers 1, 4 and 5 correspond to overrepresented sequences in FastQC reports generated from fastq files containing reads mapped exclusively to the SARS-CoV-2 genome; primers 2, 3, 6 and 7 correspond to reads showing high coverage in .BAM files. Primers 8 - 10 are negative controls that anneal to regions with low coverage in .BAM files. Data were normalized to endogenous miRNA let-7a-5p.

## RESULTS

### Small RNAs derived from SARS-CoV-1 and SARS-CoV-2

To explore the repertoire of short viral RNAs, we analyzed publicly available small RNA-seq datasets from Calu-3 cells infected with SARS-CoV1 or SARS-CoV2 (**Fig.1A**) [Wyler et al, 2021], which were, to our knowledge, the only available raw sequencing data generated by a library preparation kit not biased towards 5’-Phosphate/3’-OH small RNAs. We reasoned that viral short RNA fragments might be used for COVID-19 diagnosis and monitoring using qPCR-based assays. For this purpose, small RNA-seq reads from both mock and infected cells were trimmed from adapters, size-selected and aligned to the corresponding viral reference genome. Next, the count-per-million (CPM) values of reads mapped to each reference were calculated. Infection with either virus caused robust accumulation of short RNA fragments derived from both forward and reverse strands of viral RNA genomes 4, 12 and 24h post infection in Calu-3 cells (**Fig.1B**). Furthermore, the number of reads mapped to the forward viral RNA strands was approximately 20-fold higher than those aligned to the reverse strands (**Fig.1B**), perhaps due to greater protection of (+) genome strands by ribosomes or other associating proteins. Perfect-match viral sequences in the small RNA fractions of Calu-3 cells 24 h post-infection were 2.83% (2.68% forward + 0.15% reverse) of the total for SARS-CoV-1 and 1.51% (1.42% forward + 0.09% reverse) for SARS-CoV-2 (**Fig.1C**). This almost 2-fold difference could be explained by significantly higher cytotoxicity of SARS-CoV-2 [Wyler et al, 2021]. As could be expected, some SARS-CoV-1 small RNA reads aligned to the SARS-CoV-2 reference genome and *vice versa* due to the inherent similarity of the genomes (**Fig.1C**).

**Figure 1.**
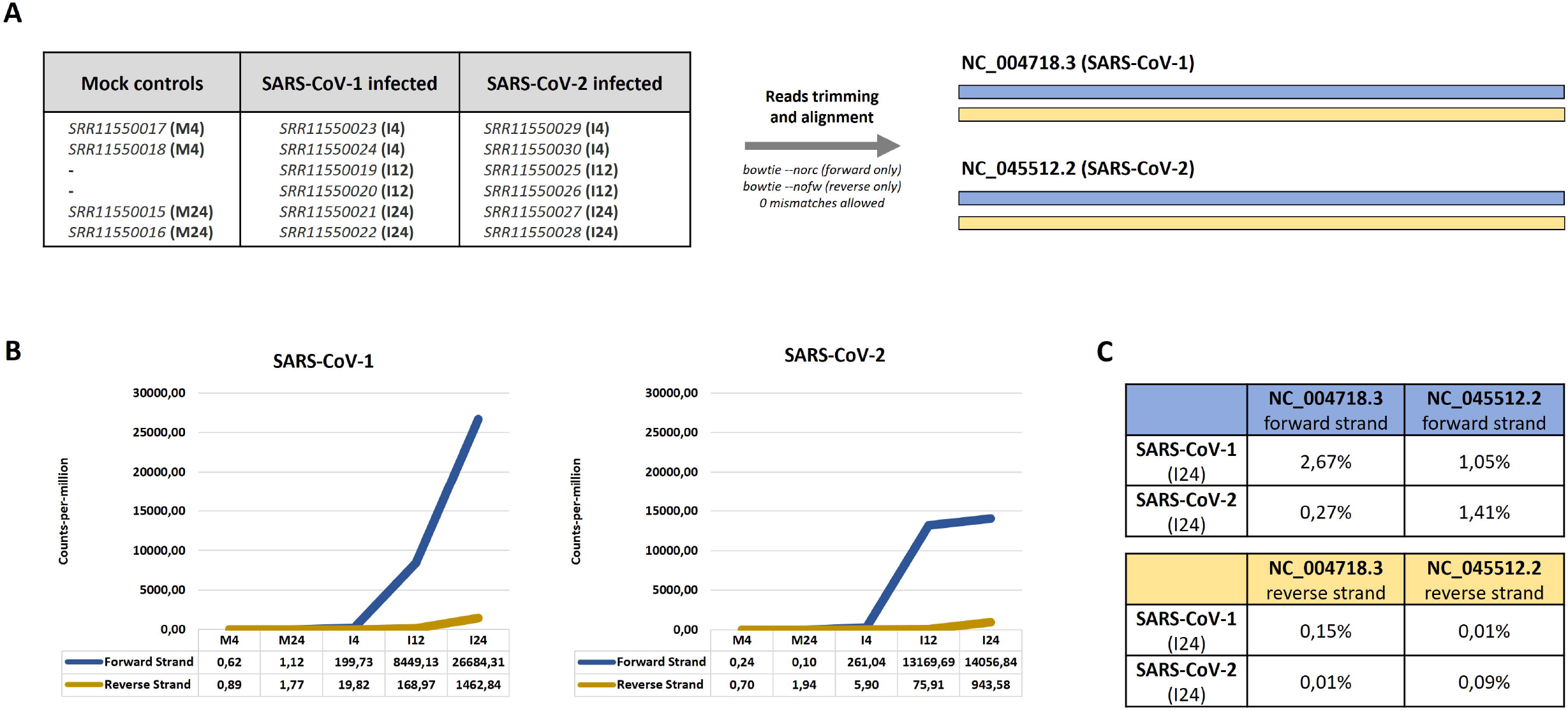
Overexpression of SARS-CoV-1 and SARS-CoV-2 derived short viral RNAs in infected Calu3 cells. **(A)** Accession numbers of small RNA-seq datasets from [Wyler et al, 2021] and the parameters used for reads alignment to corresponding viral genome references. **(B)** Expression of viral small RNA reads (in count-per-million metrics) in Calu3 cells infected with either SARS-CoV-1 or SARS-CoV-2 at 4 hours (I4), 12 hours (I12) and 24 hours (I24) post-infection. The viral RNA content in mock (uninfected) cells at 4 hours (M4) and 24 hours (M24) are also shown; **(C)** Percentage of total small RNA-seq reads aligned to forward and reverse strands of SARS-CoV-1 and SARS-CoV-2 genomes calculated at 24 hours post-infection. Note: a certain percentage of reads from SARS-CoV-1 infected cells aligned to the SARS-CoV-2 genome and *vice versa*.

### Dissection and validation of SARS-CoV-1/2-derived short RNAs

Next, we aimed to evaluate the distribution pattern of viral short RNA (vsRNA) reads and to identify virus-specific fragments mapping to discrete locations (peaks). For this purpose, we extracted reads aligning exclusively to either the SARS-CoV-1 or SARS-CoV-2 genome (forward or reverse strands), discarding overlapping fragments. As evident from the genome browser maps, multiple reads aligned discretely to single positions within SARS-CoV-1 or SARS-CoV-2 genomes (**Fig.2, Supplementary Fig. 1**, **Supplementary Fig. 2**).

**Figure 2.**
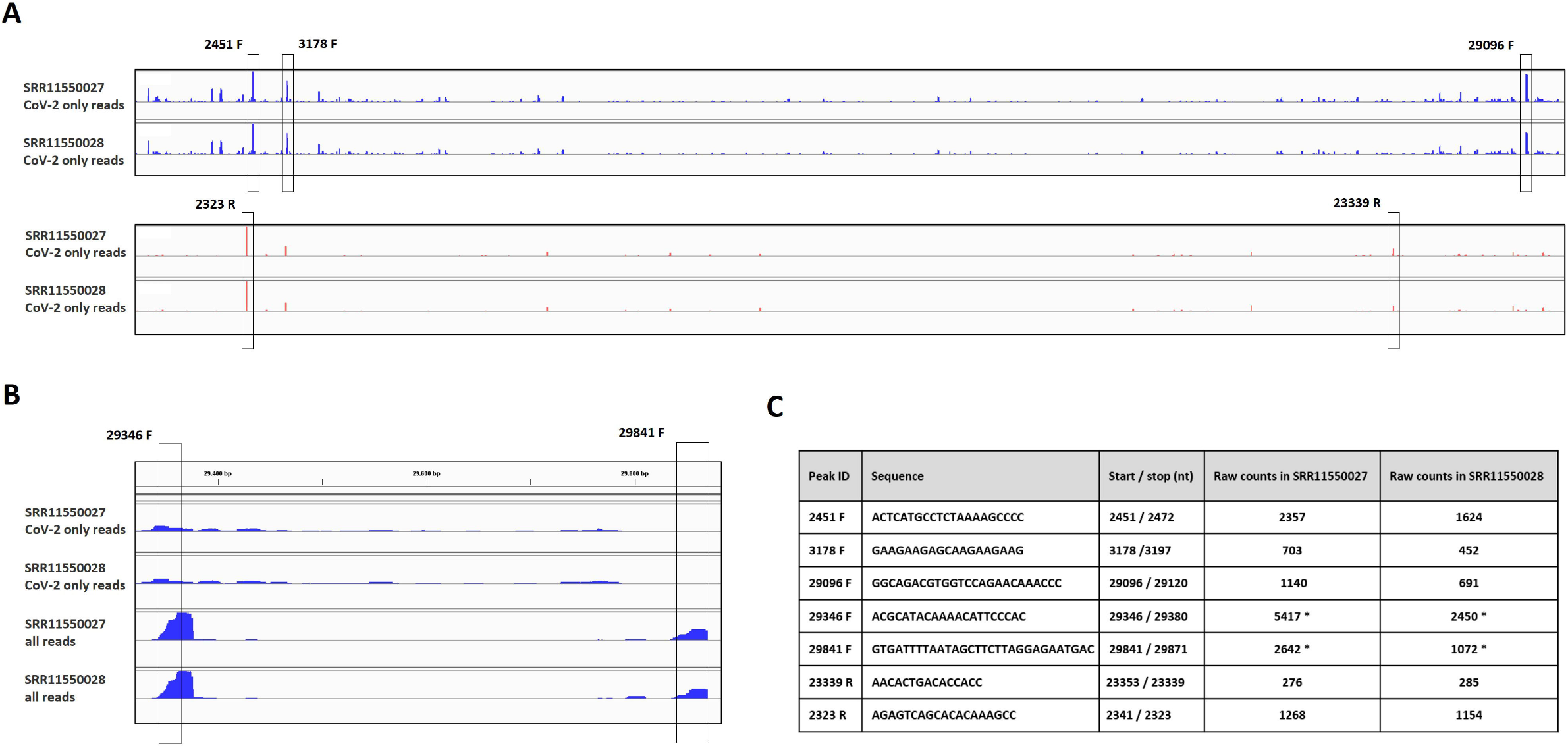
Short RNAs derived from SARS-CoV-2. **(A)** Genomic IGV map of reads aligned to the SARS-CoV-2 genome only (overlapping SARS-CoV-1 reads were filtered) generated from small RNA-seq datasets of Calu3 cells 24 hours post infection. The five most overrepresented short viral RNAs (deriving from forward and reverse strands) are highlighted by vertical rectangles; **(B)** Genomic IVG map showing the regions with highest coverage towards two motif sequences from total reads and SARS-CoV-2 reads only. **(C)** List of highly overrepresented viral sequences selected for RT-qPCR validation and their copy numbers in small RNA-seq datasets from Calu3 cells 24 hours post-infection with SARS-CoV-2. (*) – the number of times a given sequence is present in the corresponding dataset as counted using the *grep* command in Unix Bash/Shell.

To assess the presence of five selected viral short RNAs annotated as 2451F, 3178F, 29096F, 23339R and 2323R, along with two RNA sequences with high local coverage - 29364F and 29841F (**Fig.2**), in a different infection setting, we used RNA from infected and uninfected A549 ACE2-cells (**Fig.3**). Polyadenylation and reverse transcription were followed by real-time quantitative PCR (**Fig.3A, Fig.3B**). For all five putative forward-strand svRNAs (2451F, 3178F, 29096F, 29364F and 29841F), signal in infected cells was greatly increased over background (**Fig.3C**). In contrast, reverse-strand fragments 23339R and 2323R were not detected over background (**Fig.3C**). Importantly, dependance on the addition of poly(A) tails to the 3’ end of small RNAs supports the notion that detected vsRNAs were amplified from bona-fide small RNA templates rather than full-length viral RNAs. We also tested three negative control primers that annealed to SARS-CoV-2 genomic regions with fewer than 30 counts in the coverage plots (**Fig.3C**). For two, signal was similar in infected and uninfected cells (**Fig.3C**). Interestingly, the third negative control (nc14008) appeared to be induced in infected cells. However, amplification occurred even later than for the putative SARS-CoV-2 small RNA with the latest amplification (Cq 25.5 vs 23.0) (**Fig. 3C**). These data thus suggest that the SARS-CoV-2 genome may encode small RNA fragments of a discrete size.

**Figure 3.**
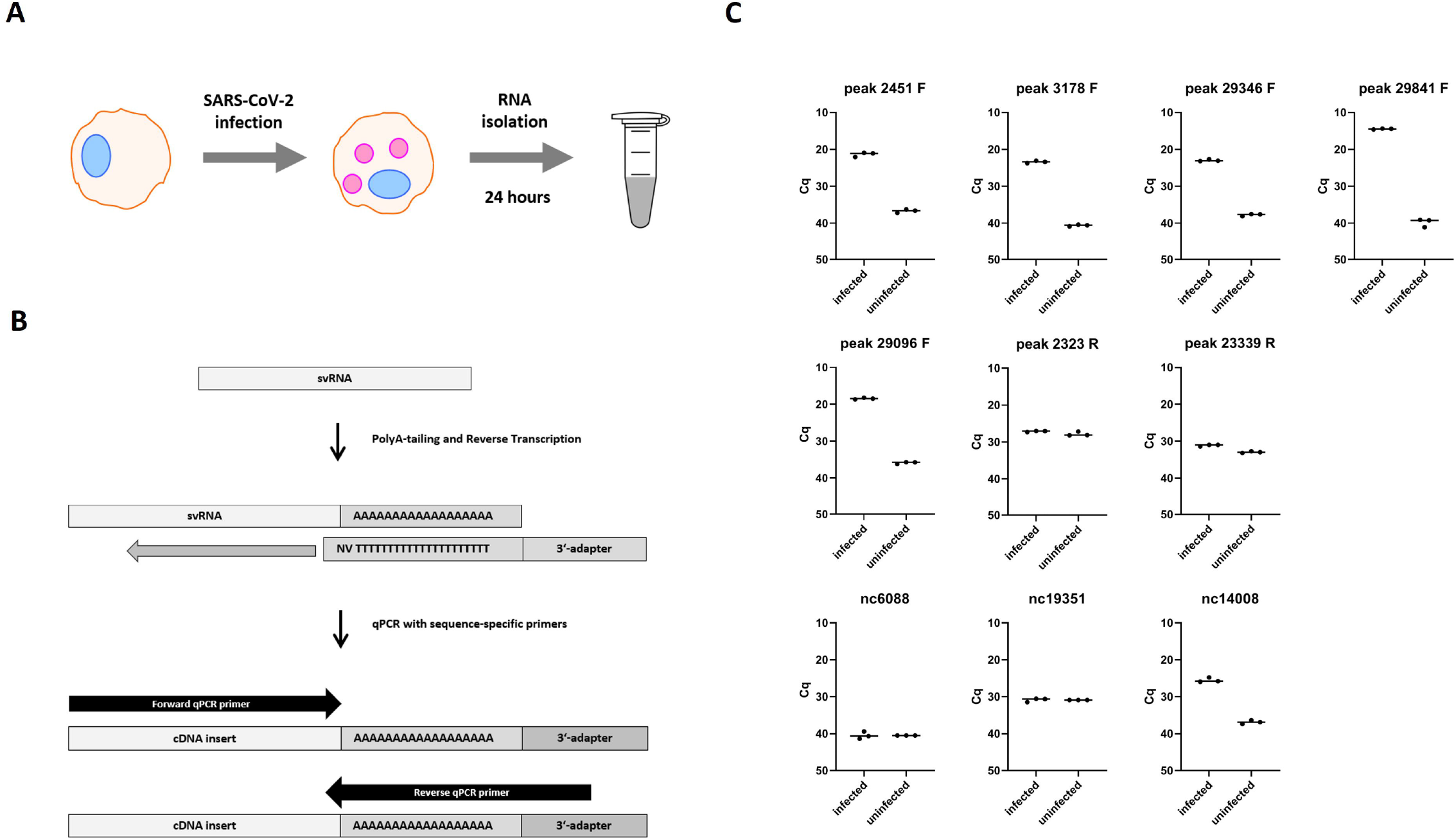
Experimental validation of several identified svRNAs with independently infected A549 cells. Experimental workflow **(A)** and RT-qPCR strategy used for short RNAs detection **(B)**. **(C)** Graphs showing Cq values for the analyzed short viral RNAs in uninfected vs. infected A549 cells.

### Read counts of SARS-CoV-2 viral miRNAs versus other short RNAs

Several groups have previously reported putative SARS-CoV-2-derived endogenous miRNAs [Pawlica et al, 2021; Singh et al, 2022; Zhao et al, 2022]. Specifically, using deep sequencing of small RNAs carrying 5’-phosphate and 3’-OH isolated from infected Calu-3, A549 and PC-9 cells, Pawlica and co-authors identified “CoV2-miR-O7a,” encoded in the SARS-CoV-2 ORF7 [Pawlica et al, 2021]. These results were confirmed by Singh et al, who detected CoV2-miR-O7a.1 and its isoform CoV2-miR-O7a.2 in infected Caco-2 and A549-ACE2 cells [Singh et al, 2022]. Finally, CvmiR-5 was identified as a viral miRNA encoded in ORF1a [Zhao et al, 2022]. In the *SRR11550027 and SRR11550028* datasets (small RNA-seq of cells 24h post-infection) only a small number of these viral miRNA sequences were found. Specifically, the top three short viral RNAs identified in our study were up to 100-1000-fold more abundant than previously reported viral miRNA reads in the same samples (**Table 2, Fig.2C**).

**Table 2.**
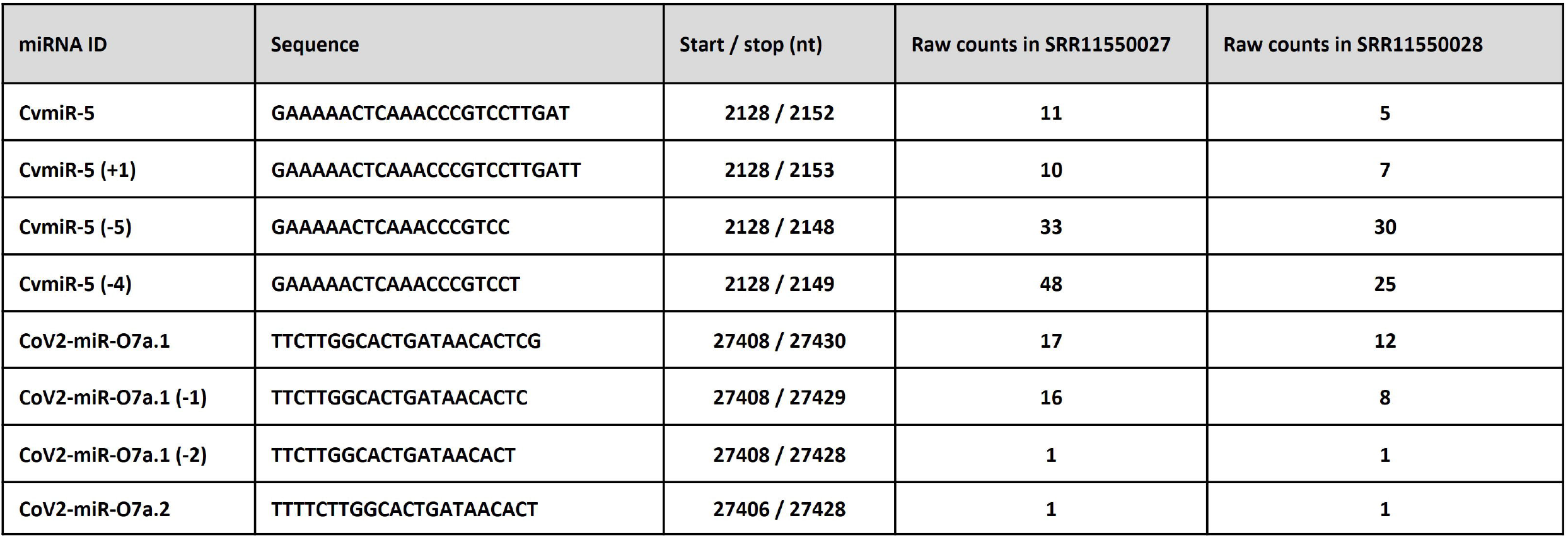
**List of previously reported miRNAs encoded in SARS-CoV-2 genome** and their copy number in SRR11550027 and SRR11550028 small RNA-seq datasets from [Wyler et al, 2021] counted using Bash/Shell *grep* command.

### Host small RNA expression in control and infected cells

In their original publication, Wyler et al restricted differential expression analysis of small RNA-seq data in control and infected cells to miRNA as well as vault RNA [Wyler et al, 2022]. We extended the analysis of their datasets to the whole-transcriptome level by mapping total reads to a custom-curated human reference transcriptome in a sequential manner as described in Materials and Methods (**Fig.4A**). Subsequently, differential expression analysis was performed for mock-infected controls versus SARS-CoV-1 and SARS-CoV-2-infected cells at 24h post-infection. Significantly more small RNA transcripts were differentially expressed in SARS-CoV-2 versus SARS-CoV-1 infection (**Fig.4B**). Specifically, of 14747 transcripts remaining after low-expression filtering, only 12 up- and 0 down-regulated sequences were identified in SARS-CoV-1 infection, compared with 268 up- and 120 down-regulated sequences in SARS-CoV-2 infection (**Fig.4C**). Only 7 differentially expressed transcripts overlapped between SARS-CoV-1 and SARS-CoV-2 infected cells (**Fig. 4B**). Interestingly, the percentage of reads mapped to mature miRNAs and pre-miRNAs combined was significantly lower than the proportion of reads mapped to other RNA classes (**Fig. 4D**). Thus, only 1 and 45 miRNAs/premiRNAs showed significant (adj. p-values < 0.05) differential expression in SARS-CoV-1 and SARS-CoV-2 infection, respectively. While hsa-miR-155-3p was markedly upregulated in both SARS-CoV-1 and SARS-CoV-2 treated cells (LFC 4.0 and 2.7, respectively), the majority of miRNAs/premiRNAs showed relatively subtle changes 24h after SARS-CoV-2 infection (−2 < LFC < 2). In contrast, fragments of protein-coding mRNAs and lncRNAs predominated among the differentially expressed transcripts (**Fig. 4D, Supplementary Table 1**). Finally, SARS-CoV-2 infection induced a strong upregulation of short RNA fragments deriving from 3’- and 5’-termini of fulllength Y-RNAs (LFC 3.0), and to a lesser extent vault RNA (LFC 1.61), as well as tRNA (LFC 1.25) (**Fig. 4D**). On the levels of individual tRNAs, however, a more profound upregulation was observed, with LFC values between 2.94 - 5.06 for the top 10 differentially expressed tRNAs (**Supplementary Fig. 3**). Interestingly, the size distribution of reads mapped to Y-RNAs, vault RNAs and tRNAs was noticeably different in SARS-CoV-2 infected cells as compared with mock-infected controls, indicating that the infection induced generation of a distinct set of short RNAs from the aforementioned sncRNAs (**Supplementary Fig. 4**). The full-length Y-RNAs, vault RNAs and tRNAs were not found in the small RNA-seq dataset, indicating that the methods applied for RNA isolation and library preparation were highly biased to capturing RNAs of shorter sizes (**Supplementary Fig. 4**). In addition, RNY, vault RNA and tRNA reads mapped predominantly to 3’- and 5’-termini of the parental transcripts (**Supplementary Fig. 5-7**). Finally, a number of miscellaneous Y-RNA transcripts, mapped to various Y-RNA pseudogenes scattered across human genome, were found in the small RNA datasets from SARS-CoV-2 infected cells (**Supplementary Fig. 3, Supplementary Fig. 8).**Many of these minor Y-RNA sequences have very marginal presence in the uninfected cells but accumulated robustly within 24h of SARS-CoV-2 infection (**Supplementary Fig. 3).**

**Figure 4.**
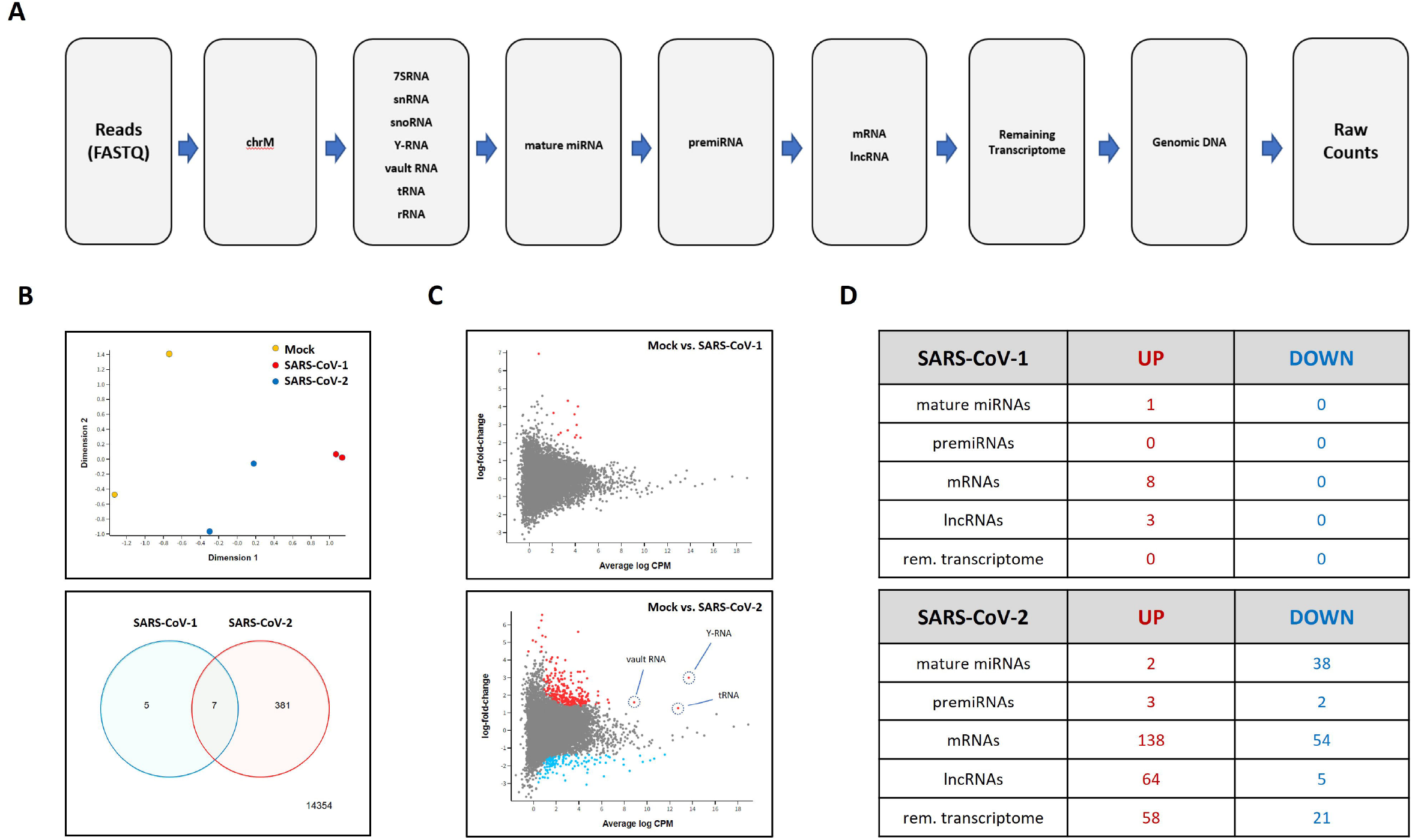
Small RNA-seq data analysis of Calu-3 cells 24-hour post-infection with SARS-CoV-1 and SARS-CoV-2. **(A)** Sequential reads mapping workflow used to generate count tables from raw small RNA-seq data; **(B)** *Upper*: Multidimensional scaling (MDS) plot of log-CPM values over dimensions 1 and 2 with samples colored and labeled by sample showing distances between the compared groups of samples; *Lower*: Venn diagram showing the number of DE genes in the comparison between SARS-CoV-1 infected cells vs. mock control (left), SARS-CoV-2 infected cells vs. mock control (right), and the number of genes that are DE in both comparisons (center) The number of genes that are not DE in either comparison are marked in the bottom-right; **(C)** Mean-difference (MD) plots of differentially expressed genes at small RNA level between mock and SARS-CoV-1 (*upper*) as well as mock vs. SARS-CoV-2 (*lower*) infected Calu-3 cells. Significantly upregulated (red dots) or downregulated (blue dots) genes were defined as those with a Benjamini-Hochberg-corrected p value < 0.05. **(D)** Table summarizing the number of differentially expressed transcripts for each RNA biotype.

### Short RNAs deriving from SARS-CoV-1/2 also align to various hg38 sequences

Next, we explored whether identified SARS-CoV-1 and SARS-CoV-2 viral short RNAs have similarities with human sequences, and thus could potentially target host transcripts. For this purpose, we aligned viral RNA reads extracted from the investigated small RNA datasets to human transcriptome references and subsequently genomic DNA with 0 mismatch tolerance (**Fig. 5A**). Mapping was done to forward (*botwie* --*norc*) and reverse (*bowtie* --*nofw*) orientations separately, identifying viral RNAs aligning to hg38 transcriptome in both sense and antisense orientation (**Fig. 5A**). The reads that did not align to human transcriptome references were further mapped to human genomic DNA (without strand specificity), which allowed us to separate reads mapped to introns and intergenic regions exclusively. Interestingly, 27.23% - 30.87% (depending on mapping) of total SARS-CoV-1 and 16.49%- 16.81% of total SARS-CoV-2 reads aligned to human introns and intergenic regions (**Fig. 5B**). Furthermore, a significant proportion of total viral reads were mapped to human protein-coding and lncRNA transcriptome. It should be noted that the vast majority of SARS-CoV-1 and SARS-CoV-2 reads that aligned to human sequences were 15-18 nt in size (**Fig. 5C, 5D**) and mapped to a single location within certain mRNA or lncRNA genes.

**Figure 5.**
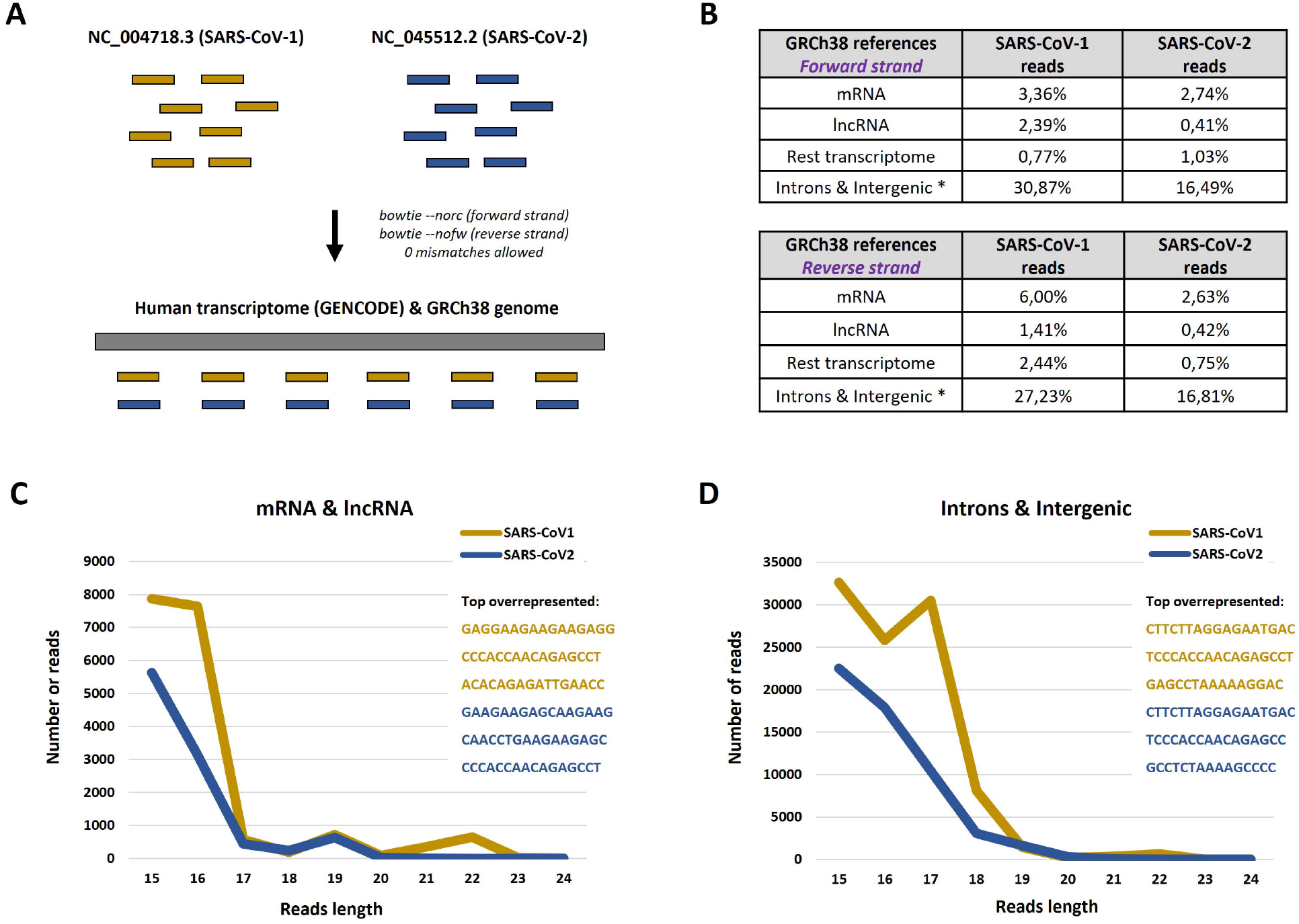
Alignment of short viral RNA reads to human sequences. **(A)** RNA-seq read mapping workflow and parameters. **(B)** Percentage of total viral small RNA reads (from both forward and reverse strands) aligned to certain hg38 sequences in forward (command: *bowtie* --*norc*) and reverse (command: *bowtie* --*nofw*) orientations. (C) Length distribution of SARS-CoV-1 and SARS-CoV-2 reads mapped to human mRNA and lncRNA references (combined). (D) Distribution of length of SARS-CoV-1 and SARS-CoV-2 reads mapped to human introns and intergenic regions (combined).

## DISCUSSION

The genomes of SARS-CoV-1 and SARS-CoV-2 are positive-sense single-stranded RNAs that utilize translational machinery of the host cell to produce proteins necessary for replication of viral RNA and assembly of viral particles [Fehr and Perlman, 2015; Naqvi et al, 2020, Kim et al, 2020]. The host cell detects and counters the pathogen through pathways of the innate antiviral immune response such as dsRNA-mediated activation of Toll-like receptors [Lowery et al, 2021, Ricci et al, 2021, Zhao et al, 2021]. SARS-CoV-1 and SARS-CoV-2 replication both involve synthesis of various RNA intermediates [Kim et al, 2020; Zhao et al, 2021; Bruce et al, 2022] whose stability and diagnostic potential remains largely unexplored. Current PCR-based methods for COVID-19 diagnostics rely on certain amplicons within the full-length viral RNA.

The original goal of this work was to assess the length and distribution of small RNAs derived from separate SARS-CoV-1 and SARS-CoV-2 genome strands in infected human lung cells and to establish proof-of-principle that coronavirus infection can be detected by ultrashort RNA-targeted qPCR. Generation of small RNA species from viral RNA genomes has been shown for multiple pathogens [Perez et al., 2010, Parameswaran et al., 2010] including coronaviruses [Morales et al., 2017]. However, previous studies investigating small RNA transcriptomes from SARS-CoV-1 and SARS-CoV-2 infected cells utilized ligation-based library preparation protocols for deep sequencing that captured predominantly 5’-Phosporylated/3’-OH RNA molecules such as microRNA [Pawlica et al, 2021; Singh et al, 2022; Zhao et al, 2022]. In contrast, one recent publication used a ligation-independent technique [Wyler et al, 2021] but limited analysis to the Argonaute-associated small RNAs including miRNA and vault RNA. We re-analyzed these publicly available datasets and observed large amounts of various previously unreported short viral RNAs (svRNAs) produced in SARS-CoV-1 and SARS-CoV-2 infected cells.

Important, the svRNAs we found were much more abundant than previously reported viral miRNAs such as CvmiR-5 (which was detected in serum of COVID-19 patients) [Zhao et al, 2022], CoV2-miR-O7a.1, and CoV2-miR-O7a.2. In the datasets we analyzed, these viral miRNAs were 100-1000-fold lower than svRNAs derived from the sense viral RNA strand and 10-100-fold lower than svRNAs derived from the antisense strand. The much more subtle expression of virus-encoded miRNAs could result from slower maturation of Argonaute-associated RNAs as compared with other svRNAs and/or higher stability of the latter. We, therefore, hypothesize that the SARS-CoV-2 svRNAs described in this work could have higher diagnostic potential for COVID-19 compared with other circulating viral miRNAs.

For host RNAs, our analysis confirmed the observation of Wyler et al that only a few miRNAs were upregulated in infected cells. While several dozen miRNAs and premiRNAs were statistically significantly downregulated after SARS-CoV-2 infection, only a few showed 4-fold or greater changes. Such a limited response of the miRNA transcriptome was unexpected given the dramatic impact of both SARS-CoV-1 and SARS-CoV-2 on viability and the mRNA transcriptome of Calu-3 cells [Wyler et al, 2021]. Interestingly, we documented up- and down-regulation of multiple small RNAs aligned to human proteincoding mRNA and lncRNA transcripts in the infected Calu-3 cells versus mock controls. A high percentage of small RNA fragments aligned to protein-coding mRNA and lncRNA in human cells has been previously reported by several authors who applied ligation-free library preparation methods [Turchinovich et al, 2014; Meistertzheim et al, 2019]. The exact mechanism of biogenesis, the parental transcripts, and the biological role of the identified mlncasRNA remains to be elucidated. It is feasible that these RNAs could possess certain regulatory roles but might also persist as simple by-products of long RNA degradation and the background transcription level. Finally, housekeeping short non-coding RNAs including Y-RNA, tRNA and vault RNA were also upregulated 24 hours after SARS-CoV-2 infection.

The significant (up to 8-10-fold) upregulation of short RNAs deriving from all four human Y-RNA genes (RNY1, RNY3, RNY4 and RNY5) following SARS-CoV-2 infection is, to our knowledge, reported for the first time. In addition to bona fide Y-RNAs, multiple Y-RNA pseudogenes scattered across the human genome were also differentially regulated. While the number of reads deriving from Y-RNA pseudogenes constituted only a minor fraction, certain miscellaneous transcripts demonstrated more profound upregulation as compared with the four main Y-RNA transcripts. It should be noted however, that all observed RNY reads were significantly shorter than full-length Y-RNAs and mapped predominantly to 3’- and 5’-termini of the parental transcripts. One previous study demonstrated that full-length Y-RNAs might not be efficiently captured by the standard reverse transcription reaction and thus remain poorly detectable by NGS [Driedonks et al, 2018]. On the other hand, short Y-RNA-derived fragments are permissive for sequencing [Driedonks et al, 2018].

Similar experimental artifacts may also be responsible for the absence of full-length vault RNA and tRNAs in the analyzed small RNA-seq datasets. For instance, the tRNA cleavage and increased generation of tRNA-derived RNAs (tDRs) in response to cellular stress is a well-known phenomenon [Thompson et al, 2008]. However, the Tosar group has recently demonstrated that tRNA halves or tDRs can in fact be produced during RNA isolation from denaturation of full-length “nicked” tRNAs [Costa et al, 2022]. In any case, the absence of full-length tRNAs in libraries from both untreated and SARS-CoV-2-infected cells indicates that the utilized small RNA-seq kit failed to incorporate them into the final NGS library.

Interestingly, upregulation of Y-RNA, particularly RNY4, was previously described in interferon alpha (IFN-α)-stimulated B cells [Hung et al, 2015]. IFN-α is a cytokine associated with viral infection, including by SARS-CoV-2 [Galbraith et al, 2022]; therefore, increased Y-RNA levels may be triggered by IFN-α that is released during infection. Interestingly, Y-RNA is triphosphorylated and can thereby trigger cellular RNA sensors such as RIG-I, leading to IFN-α production [Hornung et al, 2006; Vabret et al, 2022]. We hypothesize that Y-RNA may play a role in a feed-forward loop driving antiviral cytokine expression during SARS-CoV-2 infection.

Finally, we found that a significant proportion of short viral RNA reads from both SARS-CoV-1 and SARS-CoV-2 also aligned to human transcriptome and genome references with perfect complementarity. Such cross-mapping could result from coronaviral sequences integration events occurring during the evolution, in a similar way as it occurred for human endogenous retroviruses [Urnovitz and Murphy, 1996]. The presence of the relatively high proportion of short RNAs derived from viral genome similar to human sequences might indicate the existence of yet uncharacterized mechanism of gene expression regulation in the host cells mediated by SARS-CoV-1 and SARS-CoV-2. At the same time, such similarity of ultra-short viral small RNAs and certain human sequences could be mainly a coincidence, without particular biological impact. However, this finding could significantly affect the results of gene expression data analysis of control and infected cells. Specifically, if the viral reads are not efficiently removed from the raw RNA-seq datasets before mapping to host genomes, multiple transcripts will manifest themselves as “upregulated” due to the crossmapping of viral RNA reads.

To summarize, in this work we show that cells infected with either SARS-CoV-1 or SARS-CoV-2 accumulate abundant previously unreported short RNAs derived mostly from the forward (+) but also the reverse (-) viral RNA strand. The fact that multiple reads mapped to single discrete locations in the viral genome suggests that these RNAs could be protected from RNase degradation by RNA-binding proteins. We hypothesize that some of these SARS-CoV-2-derived short RNAs could be used for qPCR-based diagnosis of early or persistent COVID-19 infection. The sensitivity of such qPCR assays versus conventional approaches remains to be further evaluated. While further studies are required to elucidate the mechanist of biogenesis and biological role of these short RNAs, it is plausible that SARS-CoV-2 svRNAs might participate in regulation of replication, transcription, and translation of the virus as well as cellular response to infection.

## FIGURES AND TABLES LEGENDS

**Supplementary Figure 1**: IGV maps showing reads coverage towards SARS-CoV-1 and SARS-CoV-2 genomes in small RNA-seq datasets from Calu-3 cells 24-hours post infection with each virus. Reads mapped to forward and reverse strands are shown separately.

**Supplementary Figure 2**: IGV coverage maps of reads aligned towards SARS-CoV-2 genome only (reads aligned to SARS-CoV-1 genome were discarded) in small RNA-seq datasets from Calu-3 cells 4-hours, 12-hours and 24-hours post infection with SARS-CoV-2. Reads mapped to forward and reverse strands are shown separately.

**Supplementary Figure 3**: Raw read counts and log-fold-change values for the four major Y-RNA transcripts, as well as the top 10 miscellaneous Y-RNA and tRNAs for Calu3 cells mock infected and at 24-hours post-infection with SARS-CoV-1 and SARS-CoV-2.

**Supplementary Figure 4**: Distribution of length of the reads mapped to RNY **(A)**, vault RNA **(B)**and tRNA **(C)**in small RNA-seq datasets (*SRR11550027 and SRR11550028)* from Calu-3 cells 24-hour post-infection with SARS-CoV-2. The top 5 overrepresented reads are shown for each RNA biotype.

**Supplementary Figure 5**: IGV coverage maps of reads aligned to the main human RNY transcripts in small RNA-seq datasets from Calu3 cells mock infected and at 24-hours postinfection with SARS-CoV-2.

**Supplementary Figure 6**: IGV coverage maps of reads aligned to human vault RNA transcripts in small RNA-seq datasets from Calu-3 cells 24-hours post infection with SARS-CoV-2.

**Supplementary Figure 7**: IGV coverage maps of reads aligned to the top four upregulated tRNA transcripts in small RNA-seq datasets from Calu3 cells mock infected and at 24-hours post-infection with SARS-CoV-2.

**Supplementary Figure 8**: Distribution of length of the reads mapped to miscellaneous RNY references from GENCODE v.41 and the top 5 overrepresented reads. All reads mapped to the main four RNY genes were removed before aligning to GENCODE v.41 RNY references.

**Supplementary Table 1.xls: List of differentially expressed human transcripts in small RNA-seq datasets.**Datasets used: Calu3 cells mock infected (SRR11550015, SRR11550016) and at 24-hours post-infection with SARS-CoV-1 (SRR11550021, SRR11550022) and SARS-CoV-2 (SRR11550027, SRR11550028). Protein coding mRNAs are designated with a *gene_symbol* (e.g. *GAPDH*), lncRNAs are designated with the *gene_symbol*_nc (e.g. *MALAT_nc*), miscellaneous transcripts are designated with the *transcript_name* (e.g. *ENST00000511335.5*). Small non-coding RNA (except from miRNA and premiRNA) reads were combined to the corresponding transcript biotype: *tRNA*, *7SRNA*, *RNU*, *snoRNA/scaRNA*, *vault RNA*, *RNY* and *rRNA*.

## Notes

### Competing Interest Statement

The authors have declared no competing interest.

### Summary of Updates

A missing reference/link to Figure 2 has been added into the Results section. The legend for Table 2 has been updated.

