## Supplementary Fig. for "Viral and host small RNA transcriptome analysis of SARS-CoV-1 and SARS-CoV-2-infected human cells reveals novel viral short RNAs"

#### SUPPLEMENTARY FIGURES

Supplementary Figure 1

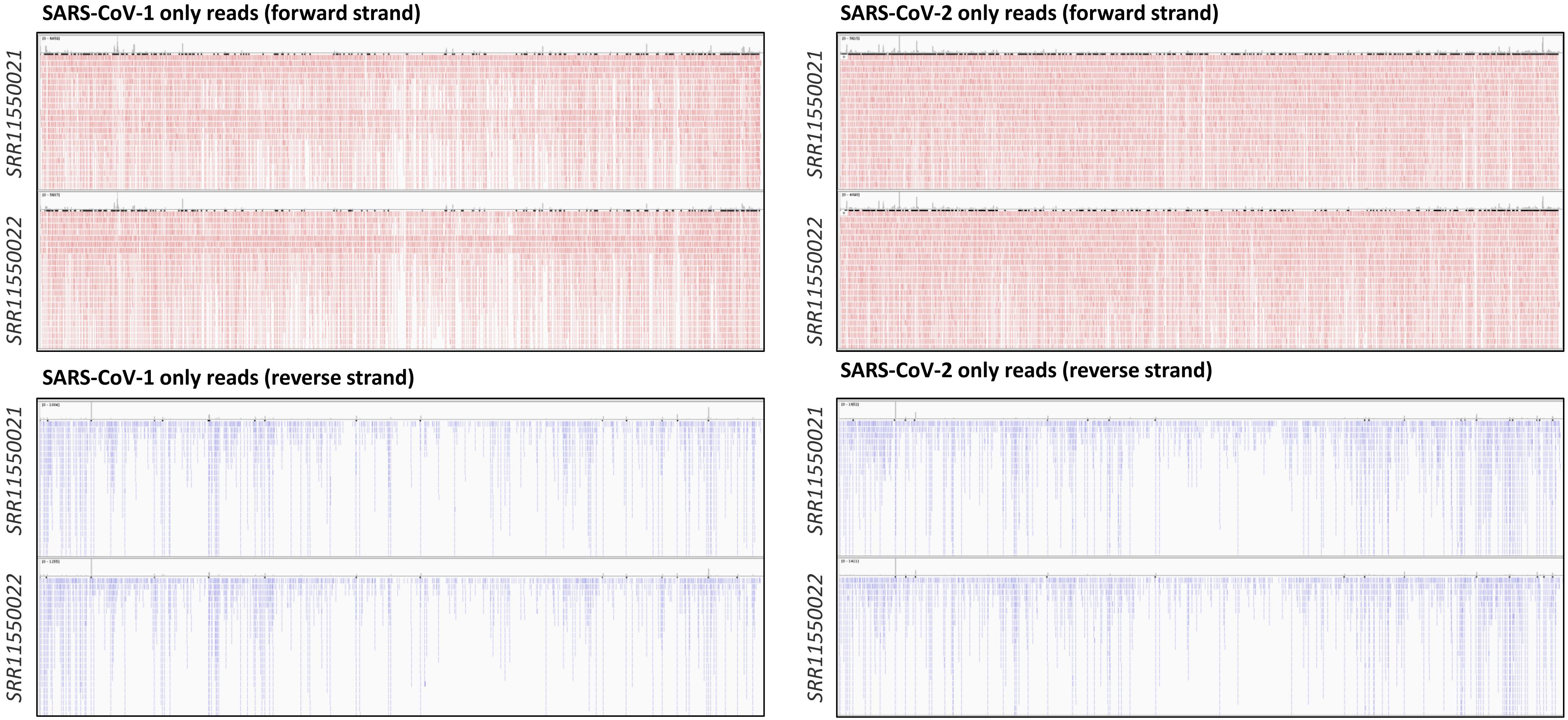

### Supplementary Figure 2

**SARS-CoV-2 only reads (forward strand)**

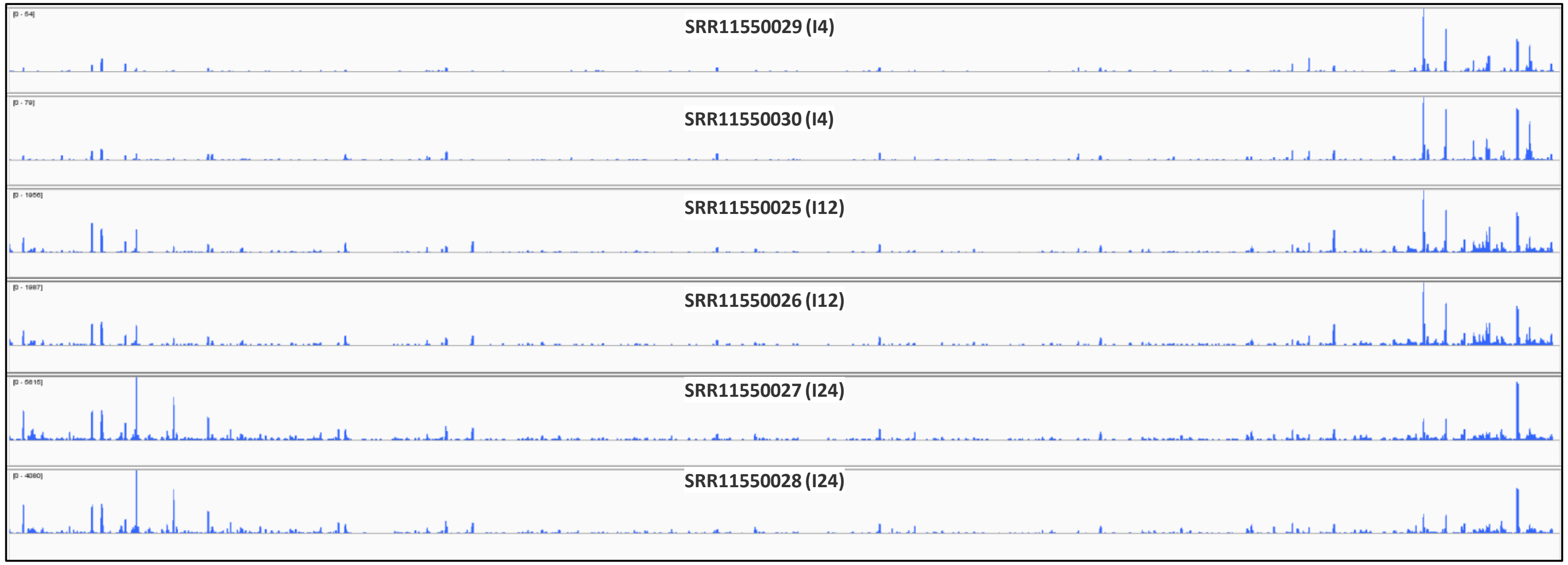


**SARS-CoV-2 only reads (reverse strand)**

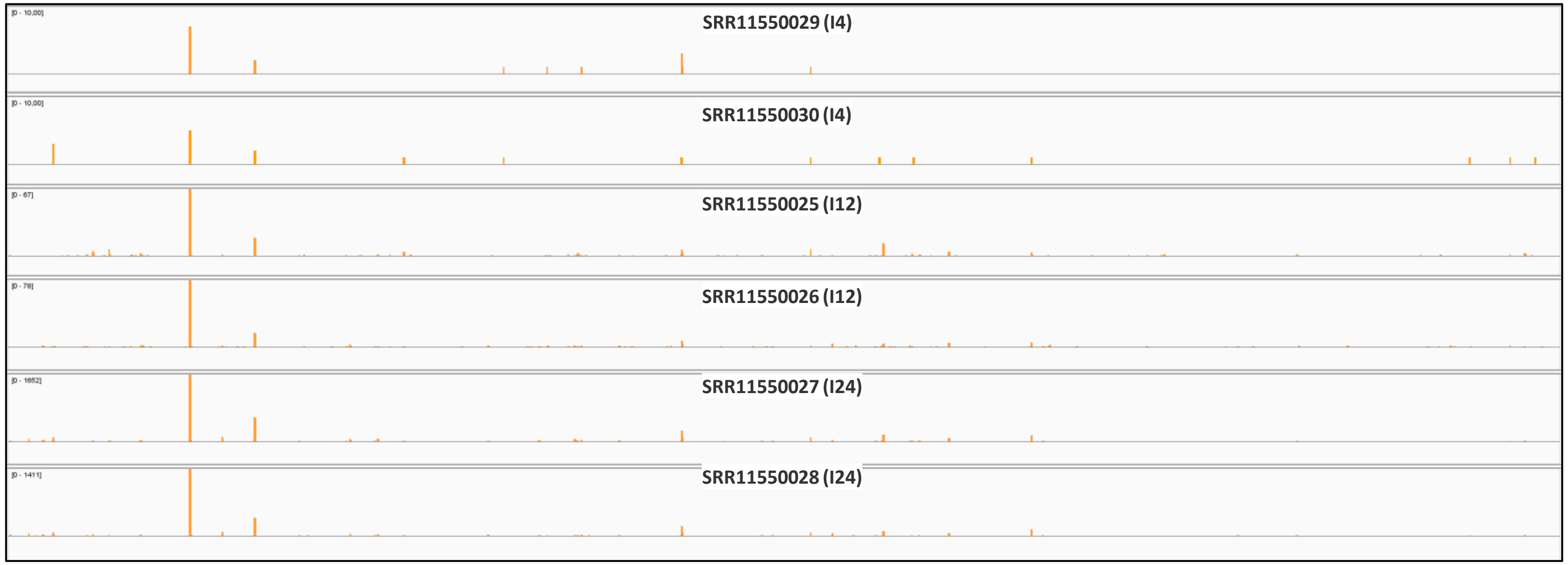

### Supplementary Figure 3

| Y-RNA *(major transcripts)* | Raw reads SRR11550015 (M24) | Raw reads SRR11550016 (M24) | Raw reads SRR11550027 (I24) | Raw reads SRR11550028 (I24) | Log-fold-change |
|---|---|---|---|---|---|
| RNY1 | 2616 | 1583 | 56859 | 26337 | 3.08 |
| RNY3 | 4688 | 2737 | 88878 | 46575 | 3.00 |
| RNY4 | 2743 | 921 | 61539 | 35294 | 3.7 |
| RNY5 | 14112 | 8914 | 158717 | 94141 | 2.28 |

| Misc. Y-RNA *(top 10 DE transcripts)* | Raw reads SRR11550015 (M24) | Raw reads SRR11550016 (M24) | Raw reads SRR11550027 (I24) | Raw reads SRR11550028 (I24) | Log-fold-change |
|---|---|---|---|---|---|
| ENST00000362540.1 | 7 | 0 | 338 | 162 | 4.74 |
| ENST00000384240.1 | 168 | 62 | 9065 | 4020 | 4.69 |
| ENST00000384001.1 | 13 | 6 | 853 | 353 | 4.53 |
| ENST00000384750.1 | 346 | 53 | 10307 | 4448 | 4.45 |
| ENST00000362354.1 | 31 | 4 | 939 | 329 | 4.17 |
| ENST00000365274.1 | 8 | 3 | 284 | 224 | 4.07 |
| ENST00000365085.1 | 11 | 1 | 264 | 164 | 3.98 |
| ENST00000459189.1 | 13 | 4 | 454 | 199 | 3.88 |
| ENST00000384478.1 | 12 | 0 | 204 | 114 | 3.79 |
| ENST00000459091.1 | 19 | 4 | 461 | 230 | 3.74 |

| tRNA *(top 10 DE transcripts)* | Raw reads SRR11550015 (M24) | Raw reads SRR11550016 (M24) | Raw reads SRR11550027 (I24) | Raw reads SRR11550028 (I24) | Log-fold-change |
|---|---|---|---|---|---|
| tRNA-SeC-TCA-1-1 | 11 | 17 | 1348 | 973 | 5,06 |
| tRNA-His-GTG-1-3 | 12 | 11 | 414 | 229 | 3,40 |
| tRNA-His-GTG-1-9 | 12 | 9 | 413 | 184 | 3,36 |
| tRNA-His-GTG-1-6 | 13 | 8 | 391 | 185 | 3,35 |
| tRNA-His-GTG-1-8 | 13 | 12 | 401 | 210 | 3,21 |
| tRNA-Lys-TTT-5-1 | 32 | 30 | 831 | 498 | 3,14 |
| tRNA-Gln-CTG-5-1 | 3 | 3 | 83 | 88 | 3,10 |
| tRNA-His-GTG-1-1 | 15 | 13 | 409 | 198 | 3,04 |
| tRNA-His-GTG-1-7 | 18 | 11 | 424 | 194 | 3,03 |
| tRNA-His-GTG-1-2 | 26 | 9 | 396 | 218 | 2,94 |

### Supplementary Figure 4

**A**

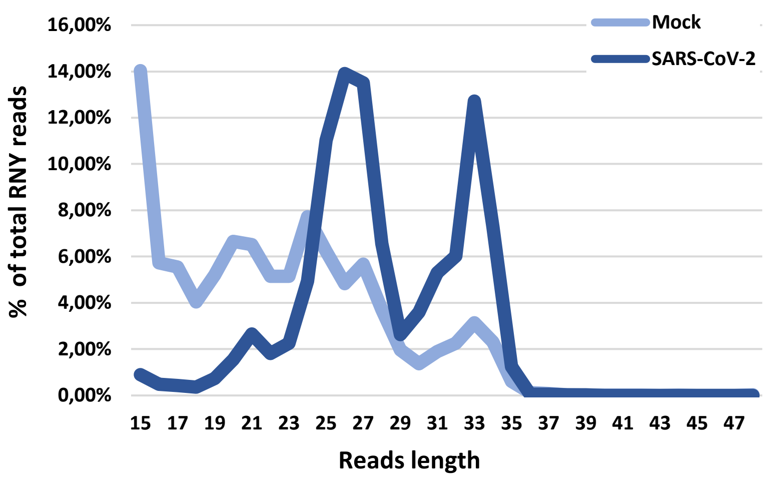


**B**

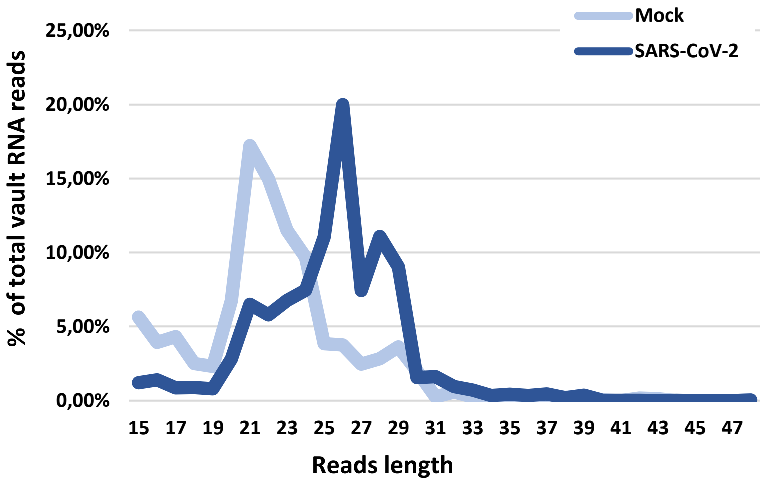


**C**

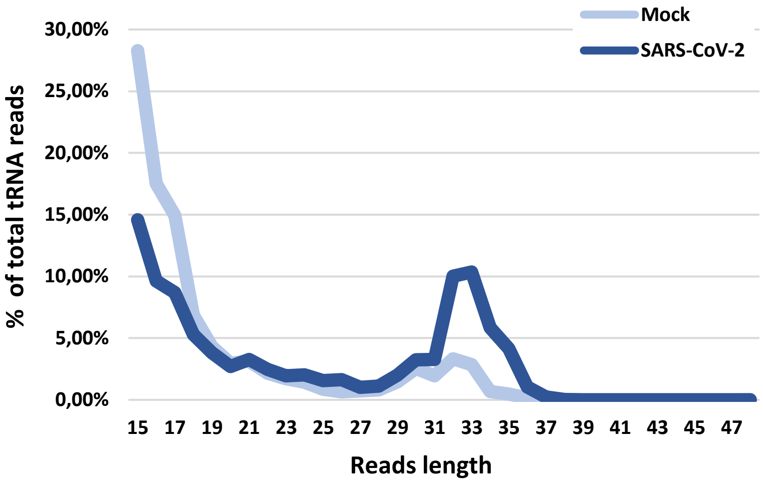


| Top 5 overrepresented RNY-mapped reads (SARS-CoV-2 infected Calu-3 cells) | % in: SRR11550027 SRR11550028 |
|---|---|
| CCCCCACAACCGCGCTTGACTAGCTTGCTGTTT | 10.5%<br>11.2% |
| CTCCCACTGCTTCACTTGACTAGCCT | 5.7%<br>5.2% |
| CTCCCACTGCTTCACTTGACTAGCC | 5.3%<br>4.7% |
| CTTCTCACTACTGCACTTGACTAGTCT | 4.4%<br>3.4%) |
| CCCCCACAACCGCGCTTGACTAGCTTGCTGTTTT | 4.4%<br>4.2% |

| Top 5 overrepresented vault RNA-mapped reads (SARS-CoV-2 infected Calu-3 cells) | % in: SRR11550027 SRR11550028 |
|---|---|
| GACCCGCGGGCGCTCTCCAGTCCTTT | 12.92%<br>11.63% |
| CGAGACCCGCGGGCGCTCTCCAGTCCTTT | 6.49%<br>4.39% |
| GAGACCCGCGGGCGCTCTCCAGTCCTTT | 5.40%<br>4.74% |
| GACCCGCGGGCGCTCTCCAGTCCTT | 4.50%<br>5.15% |
| GACCCGCGGGTGCTTTCCAGCTCTTT | 3.12%<br>2.71% |

| Top 5 overrepresented tRNA-mapped reads (SARS-CoV-2 infected Calu-3 cells) | % in: SRR11550027 SRR11550028 |
|---|---|
| GTTTCCGTAGTGTAGTGGTTATCACGTTCGCCT | 4.84%<br>2.28% |
| GTTTCCGTAGTGTAGTGGTTATCACGTTCGCC | 4.83%<br>2.22% |
| TCCCTGGTGGTCTAGTGGTTAGGATTCGGCGC | 1.63%<br>1.38% |
| GCCCGGATAGCTCAGTCGGTAGAGCATCAGACT | 1.76%<br>1.34% |
| CGAGAGGTCCCGGGTTC | 1.56%<br>2.13% |

Supplementary Figure 5

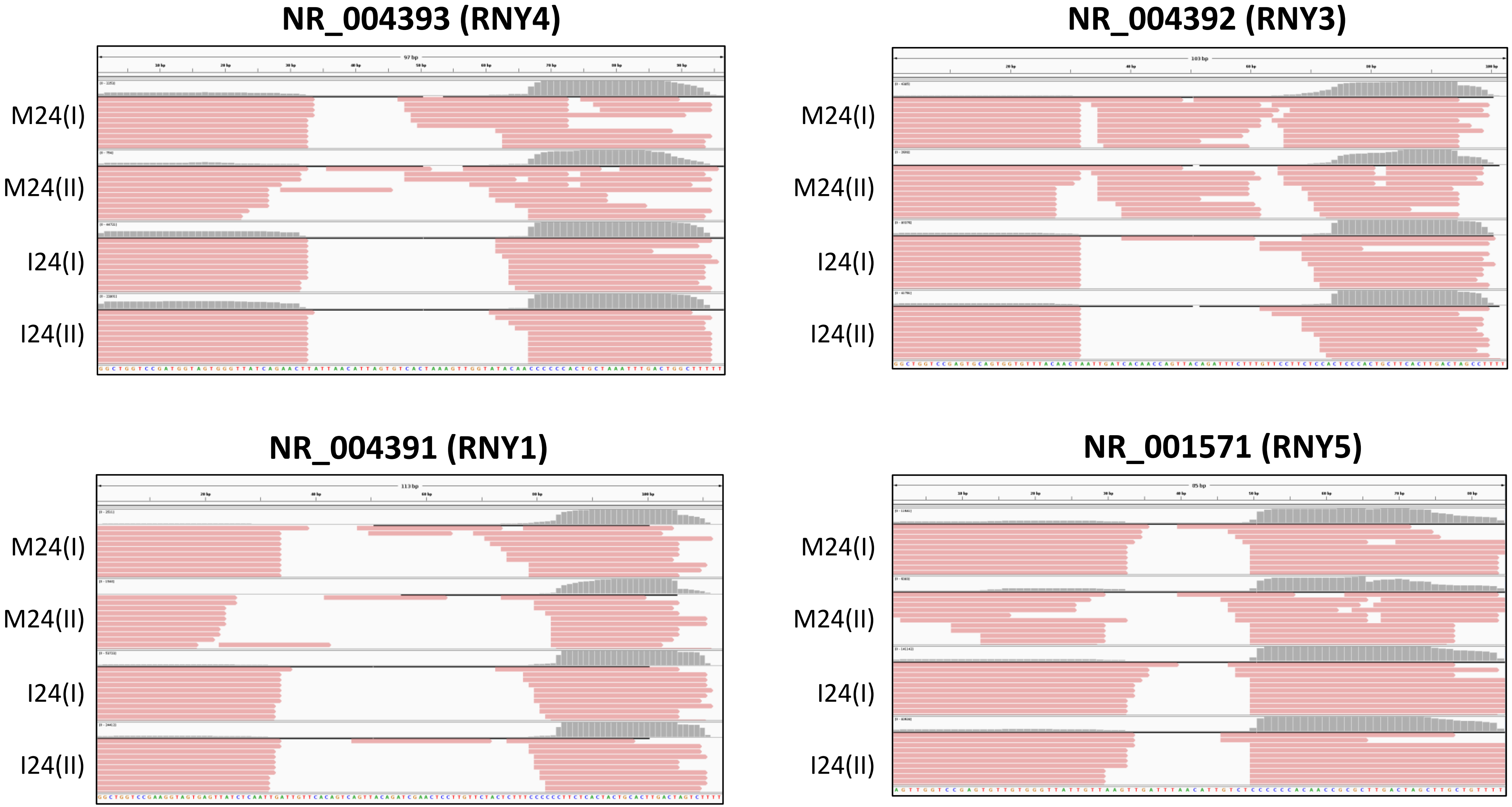

Supplementary Figure 6

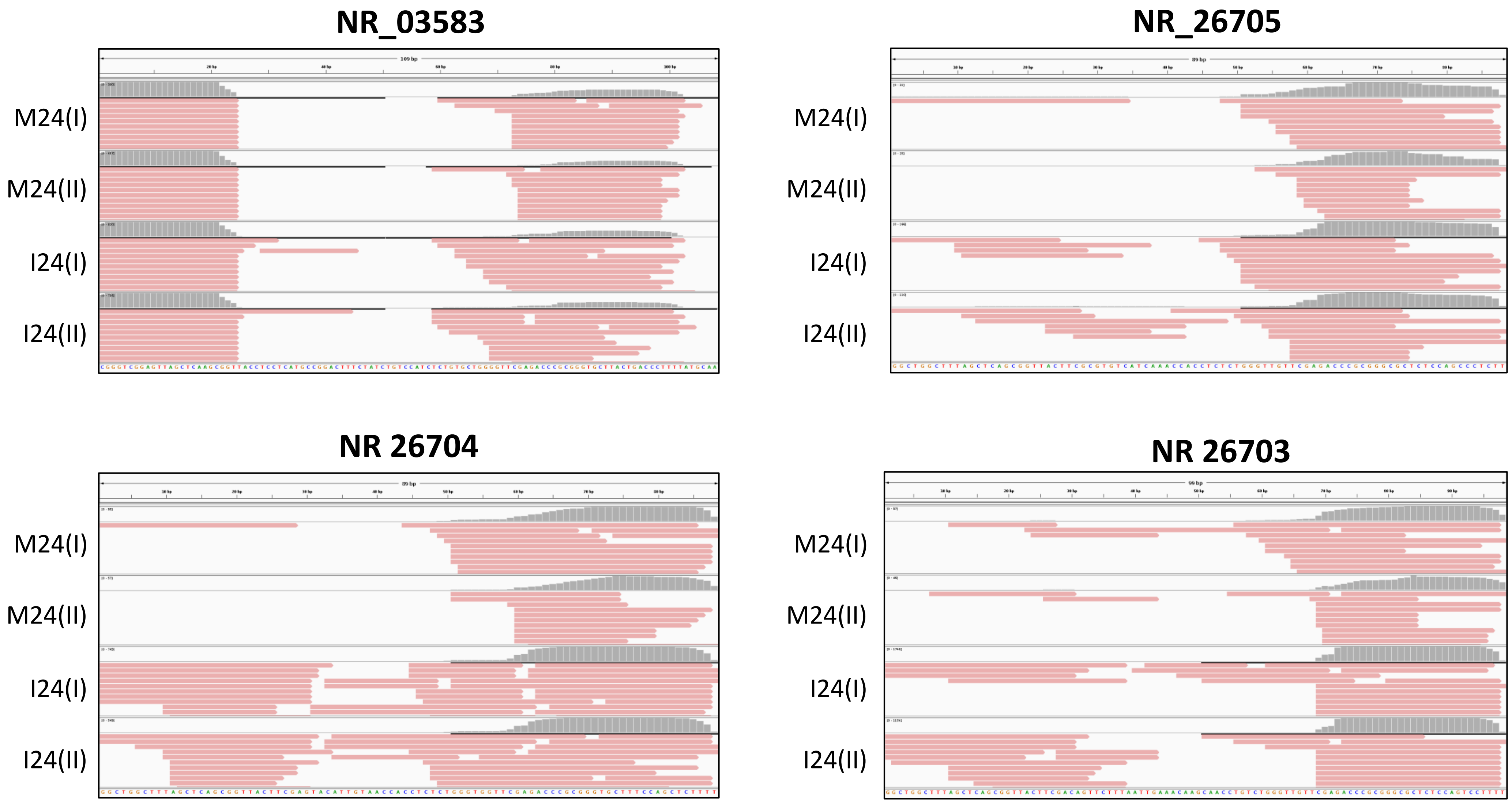

Supplementary Figure 7

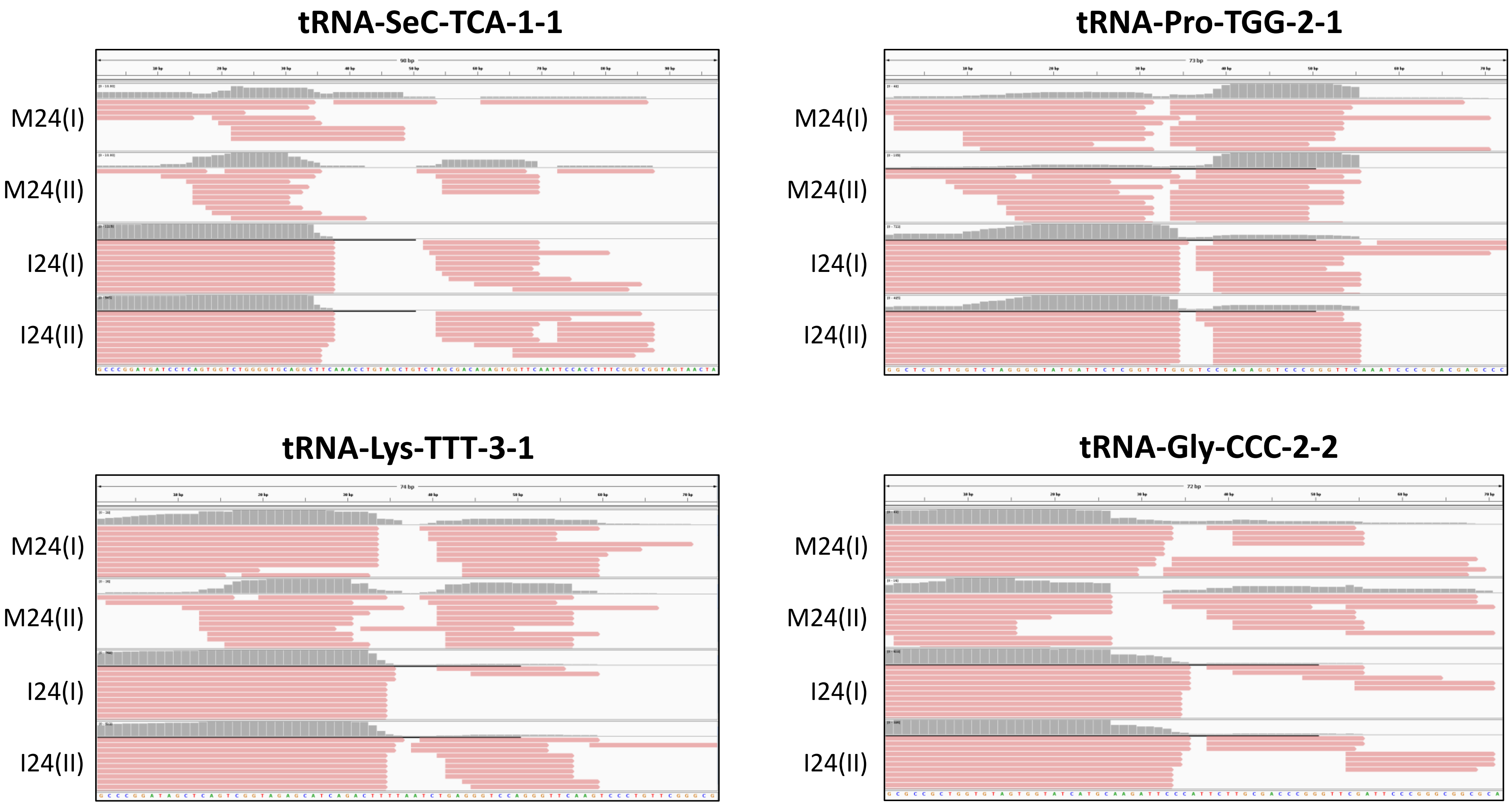

### Supplementary Figure 8

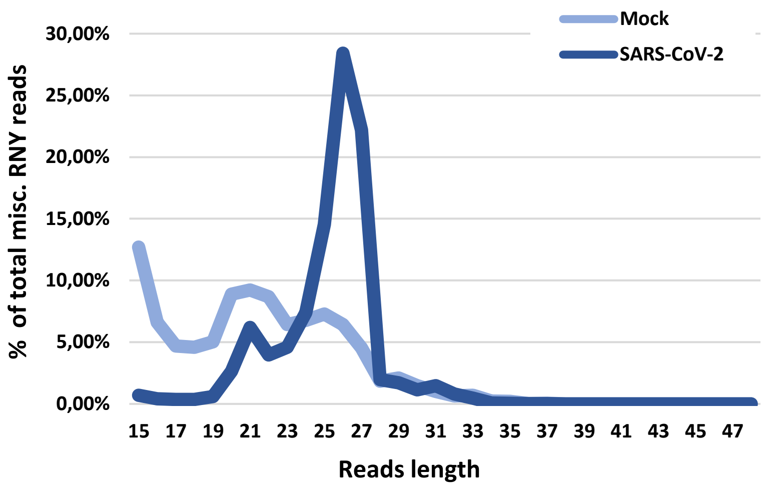


| Top 5 overrepresented misc. RNY-mapped reads (SARS-CoV-2 infected Calu-3 cells) | % in: SRR11550027 SRR11550028 |
|---|---|
| CTCCCACTGCTTCACTTGACTAGCCCT | 17.89%<br>16.64% |
| CTCCCACTGCTTCACTTGACTAGCCC | 17.38%<br>14.97% |
| TCCCACTGCTTCACTTGACTAGCCC | 4.59%<br>4.10% |
| CTTCTCACTACTGCACTTGACTAGCC | 4.25%<br>3.55% |
| TCCCACTGCTTCACTTGACTAGCCC | 3.94%<br>4.04% |
